# Windthrow-generated mounds create contrasting regeneration niches for red oak and black cherry in a deer-browsed Carolinian forest

**DOI:** 10.64898/2026.03.11.711114

**Authors:** Jiezi Duan, Kenneth A. Anyomi

## Abstract

Windstorms are a dominant recurrent natural disturbance type within Carolinian old-growth forests in southwestern Ontario. This study examined how windthrow generated microsites differ from background microsites in soil properties and seedling survival across 84 locations in Hamilton. For each site, soil samples from windthrow generated mound tops and adjacent non-disturbed forest floor were analyzed. Seedlings of red oak (*Quercus rubra*) and black cherry (*Prunus serotina*) were planted on mound tops and adjacent ground microsites, and their survival was monitored. Non-disturbed ground microsites had significantly higher soil moisture (*p* < 0.001) and organic matter (*p* < 0.001) than disturbed sites, whereas disturbed sites were consistently drier (*p* < 0.001). Red oak survival was 28% higher on the non-disturbed microsites relative to the disturbed microsites, while black cherry survival was 41% higher on disturbed microsites relative to the non-disturbed ground microsite. Logistic regression analyses reveal that soil moisture was the strongest predictor of seedling survival (*p = 0.030*), with contrasting responses between the two species (*p = 0.019*). These results suggest that windthrow generated mounds create distinct environmental conditions that selectively favor unique autecology. As windthrow frequency rises under climate change, mounds could become major mediators of forest canopy composition.

## 1.0 Introduction

Windstorms are an important recurrent natural disturbance type that shapes forest stand dynamics (e.g. Mitchel 2013). When trees are uprooted during windstorm events, the root plate lifts soil from the forest floor and forms elevated structures known as tip-up mounds, often paired with adjacent pits (Ulanova 2000). Because tip-up mounds are elevated above the surrounding forest floor, they have the potential to reduce accessibility to large herbivores such as deer, potentially creating refuges for seedling establishment (Long et al., 1998; Don et al. 2024). This is especially critical across many temperate forests where hunting restrictions and decline in natural predators have led to significant increase in ungulate browsing (Brown et al. 2000; Ballard et al. 2001). Understanding how windthrow generated mounds alter local microsites is essential in understanding potential changes in tree regeneration and forest resilience under climate change.

Tree uprooting leads to altered light environment, altered soil structure and altered soil chemistry (Schaetzl et al., 1988, Ulanova 2000, Renaudin et al. 2025). Windthrow mounds can function as localized regeneration sites by reducing competition and increasing light availability (e.g. Kern et al., 2019). A key but rarely tested assumption in forest restoration and regeneration ecology is that undisturbed forest floor microsites represent a neutral or suitable baseline for seedling establishment (e.g. Bertacchi et al. 2016). However, species differ in physiological tolerance. For example, species such as American beech (*Fagus grandifolia*), red oak (*Quercus rubra*) can persist in shaded understories through long-term advanced regeneration in compact and moist soils (Kuehne et al., 2014; Nosko et al., 2022). In contrast, species such as birch (*Betula spp*), aspen (*Populus spp*), black cherry (*Prunus serotina*) etc. are faster growing species that require well-drained soils and higher light environment (Marquis, 1990; Jagodziński et al. 2019). Understanding how recurrent windstorms impact tree regeneration and seedling establishment is essential for climate change adaptation and local restoration efforts (Bertacchi et al., 2016).

The Carolinian Forest in southwestern Ontario is a high biodiversity area that supports over 25% of the country’s at-risk species (Nature Conservancy Canada, 2025). This region also experiences the highest frequency of tornadoes in Canada (Public Safety Canada, 2020; Munoz and Anyomi, 2026), resulting in frequent windthrow events that generate mounds. At the same time, white-tailed deer (*Odocoileus virginianus*) densities in this region are estimated to be three times the ecological carrying capacity of the forest ecosystem (HCA, 2013), intensifying browsing pressure on regenerating tree species. The goal of this study was therefore to evaluate how windthrow generated microsites differ in soil properties to background (adjacent non-disturbed forest floor) microsites and their impacts on seedling survival. Specifically, we aimed to 1) Compare soil moisture, pH, organic matter, and texture between windthrow generated mound microsites and adjacent ground microsites. 2) analyze the survival of red oak (*Quercus rubra*) and black cherry (*Prunus serotina*) seedlings on mounds and adjacent ground microsites. By linking soil conditions, microsite type (mound top vs background/ground microsite), and species survival, this study aims to clarify whether windthrow generated mounds function as ecological filters that favor certain tree species and the consequences for forest composition under an intensifying disturbance regime.

## 2.0 Method

### 2.1 Study Site

This study was conducted in the old-growth Carolinian forest of Hamilton, Ontario. Forest canopy within the Hamilton township is dominated by northern temperate species such as sugar maple (*Acer saccharum*), red oak (*Quercus rubra*), American beech (*Fagus grandifolia*), shagbark hickory (*Carya ovata*), with scattered black cherry (*Prunus serotina*) and white ash (*Fraxinus americana*). Even though it occupies less than 1% of Canada’s land area, this forest type hosts 25% of Canada’s endangered species, many of which are at their northern range limit such as eastern flowering dogwood (*Cornus florida*) and American chestnut (*Castanea dentata*) among others (Pedlar et al., 2024) underscoring the importance of targeted conservation and restoration efforts.

Fieldwork was carried out in the Tiffany Falls Conservation area, McMaster Forest Nature Preserve, Borer’s Falls Conservation area, Dundas Valley Conservation area and Headwaters trail (Fig. 1).

**Fig. 1.**
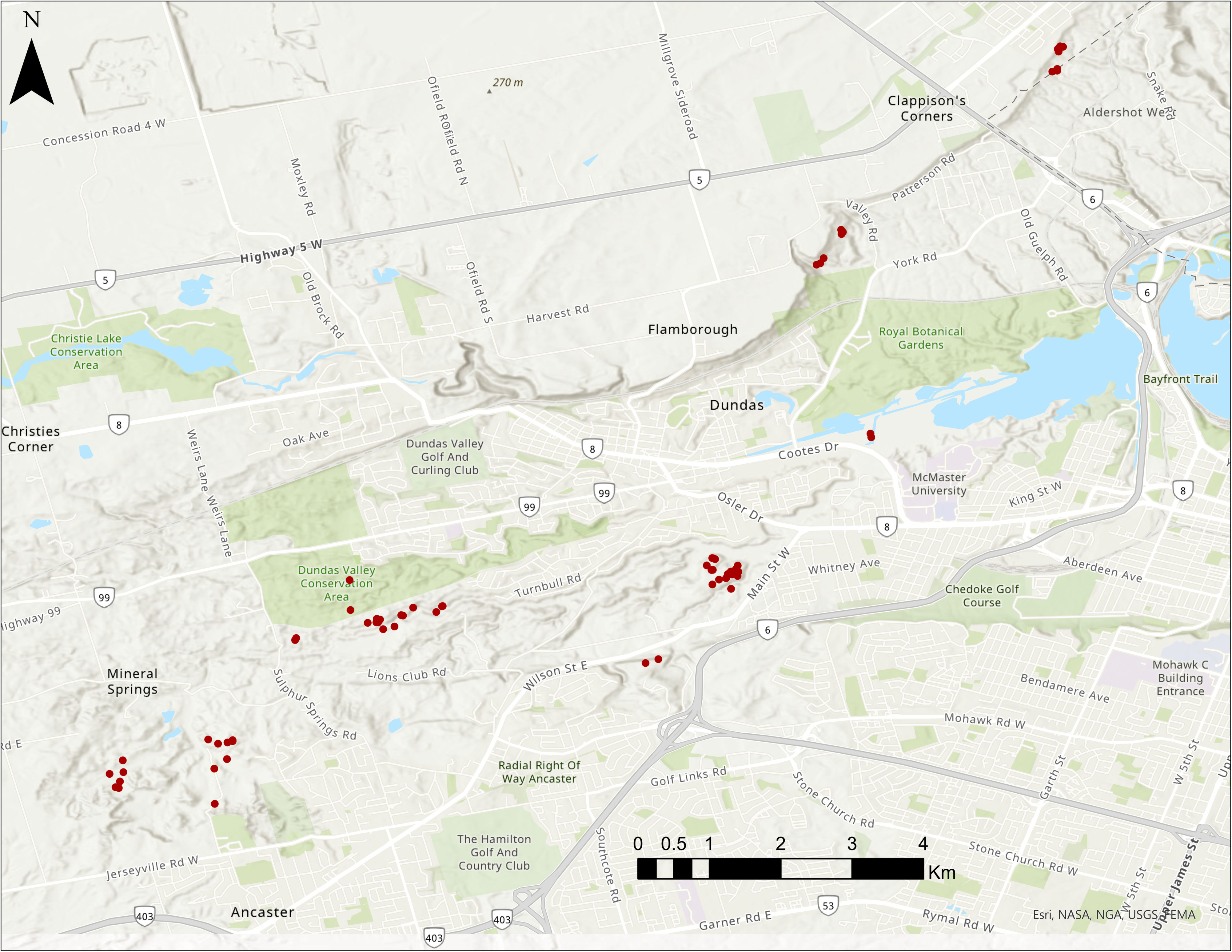
Location of windthrow mounds

### 2.2 Windthrow Mound Scouting

Field scouting was conducted to locate and assess mounds within the old-growth Carolinian forest in Hamilton which initially produced over 150 mounds and were then filtered to 84 mounds. To be considered suitable for inclusion; a) the mound had to be clearly detached from the pit, with no risk of the root plate tipping back into place - only fully uprooted and stable mounds were considered, b) the mound needed to be at least one meter in height from the base of the pit to ensure that it could potentially reduce access by deer browsing, c) there should be sufficient soil at the base of the uprooted tree to support seedling establishment. For each selected mound, GPS coordinates were recorded, and physical dimensions including mound height and width were measured.

### 2.3 Soil Sampling

For each selected mound, soil samples were collected from the windthrow mound top (disturbed site) and from the adjacent intact forest floor (ground) using a hand trowel to capture the environmental conditions directly experienced by young seedlings. Each soil sample was placed in a resealable ziplock bag, filling approximately 80 % of the bag’s volume, then sealed, labeled, and transported to the laboratory. Samples were stored in a refrigerator until subsequent analyses of soil organic matter (SOM), pH, moisture content, and texture.

### 2.4 Lab Analyses of Soil Samples

#### 2.4.1 Soil moisture and organic matter

Soil moisture content was determined gravimetrically. For each sample, approximately 100 g of fresh soil was weighed and oven-dried at 105 °C for 24 h to remove water. The dried soil mass was recorded, and the gravimetric moisture content was calculated using Eqn. 1 below.

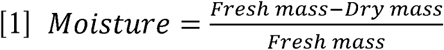

To quantify the percentage of organic matter in the soil, the dried soil samples, which contain both organic and inorganic components, were then placed into labeled ceramic crucibles and heated in a muffle furnace at 550 °C for approximately 3 hours to burn off all organic material (Hoogsteen et al. 2015). Their placement in the furnace was labelled on paper to keep track of them. After cooling for 1-2 hours, the remaining ash representing only the inorganic fraction was weighed again to obtain the ashed mass. The percentage of soil organic matter (SOM) was then calculated using the Eqn. 2 below.

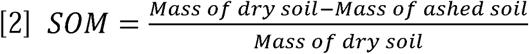

#### 2.4.2 Soil pH

Soil pH was measured using a pH probe calibrated using standard buffer solutions (pH 7.00 and pH 4.00). For each sample, 25 mL of deionized water was added to a 50 mL graduated cylinder, and soil was added gradually until the total volume reached 40 mL (≈1:1.6, v/v). Cylinders were sealed with parafilm, shaken, and allowed to settle until two layers were visible, an upper layer (water solution) and a lower soil layer. The pH probe was carefully placed into the upper water layer without disturbing the soil layer, and readings were recorded once stable. Two replicate measurements were taken for each sample, and the mean pH value was calculated.

#### 2.4.3 Soil Texture

Soil texture was determined using the mason jar sedimentation method. For each sample, a mason jar was filled to approximately half its volume with field-collected soil, after removing large stones and organic debris by hand. Deionized water was then added until the jar was nearly full, followed by approximately one tablespoon of powdered dishwasher detergent to help disperse soil particles. The contents were thoroughly mixed using a large plastic stirring rod until all soil aggregates were fully broken apart and particles were suspended in the water. The suspension was then allowed to settle undisturbed for approximately two days, until three distinct layers formed including sand (bottom), silt (middle), and clay (top). Once the layers were clearly visible, the boundaries between layers were marked on the jar. The total height of the settled material and the height of each layer were then measured with a ruler. The percentage of sand, silt, and clay was calculated by dividing each layer’s height by the total height of all layers. Soil texture class was determined by plotting these percentages on a standard USDA soil texture triangle.

### 2.5 Seedling Planting

Seedlings of two native tree species, red oak (*Quercus rubra*) and black cherry (*Prunus serotina*) which are also known to be highly preferred by deer (Barker, 2018), were sourced from a local nursery (Verbinnen’s nursery, Dundas Ontario). Red oak seedlings were approximately 20 - 40 cm in height while black cherry seedlings were 10 - 20 cm tall. For each selected mound, one seedling was planted at the top of the mound and another in the nearby intact forest floor ground microsite type, with species assigned randomly from the available stock rather than in controlled pairs. This design allowed independent evaluation of species responses to microsite conditions. On 11-very large mounds, more than one seedling was planted on top of the mound. At the time of planting, all existing leaves were removed from each seedling, leaving only the terminal bud, so that the initial leaf count for all individual seedlings was zero.

### 2.6 Monitoring Seedling Survival

Seedlings were monitored two weeks after planting (first monitoring) and at the end of the summer (second monitoring). Seedlings were recorded as dead or alive with leaves (green stem) and alive with no-leaf - both considered survivors, as both indicate the individual remained viable. Seedlings recorded as missing (absent from their planting location during monitoring) were excluded from calculations, as some may have been obscured by dense ground vegetation rather than dead or removed. Survival rate was calculated for all seedlings combined as the proportion of survivors (alive with leaves + alive with no leaf) relative to the total number of seedlings assessed (alive with leaves + alive with no leaf + dead).

### 2.7 Statistical Analysis

#### 2.7.1 Data Quality Review

Seedling survival at the end of the growing season was used as the primary response variable, as it reflected cumulative survival following planting. A total of 84 windthrow mounds were mapped and measured, each containing two microsite types i.e. mound top and adjacent ground. Although this design would normally produce 168 microsite-level observations (84 × 2), the initial dataset contained 178 observations due to two factors: (1) some mound tops contained more than one planted seedling, and (2) five ground microsites were shared by two adjacent mounds that were next to each other, resulting in one soil sample representing both mounds (a total of 5 datapoints). Data filtering was conducted at the microsite level, meaning that if either the mound top or intact forest floor ground microsite lacked valid measurements, only that observation was removed rather than excluding the entire mound. Of the initial 178 observations, 47 were excluded due to incomplete or unreliable data. These exclusions included: (1) microsites where no seedlings were planted (*n* = 12), (2) missing soil samples (*n* = 13), (3) seedlings with undetermined survival status (*n* = 4), (4) unreliable soil organic matter measurements due to container damage during loss-on-ignition (*n* = 9), and (5) invalid soil texture classifications resulting from inconsistent duplicate measurements or non-classifiable inorganic material (*n* = 9). Following these exclusions, the final dataset consisted of 131 microsite-level observations.

#### 2.7.2 Microsite Soil Attributes

Soil properties were compared by microsite type i.e. mound-top (disturbed sites) and intact ground microsites, using appropriate statistical tests based on data distribution. Welch’s *t*-test was used to compare soil moisture and soil pH by microsite types, as these variables satisfied assumptions of normality but exhibited unequal variances. Soil organic matter (SOM), which did not meet normality assumptions, was analyzed by microsite type using a Wilcoxon rank-sum test.

#### 2.7.3 Seedling Survival Analysis

Seedling survival was analyzed as a binary response variable (alive or dead at the end of the monitoring period). Survival patterns were first summarized descriptively by species and microsite to identify general trends. To evaluate the drivers of seedling survival, we fitted a generalized linear model (GLM) with a binomial error distribution and logit link function. Predictor variables included soil moisture, soil organic matter, soil pH, soil texture, species identity, and microsite type (mound top vs ground), along with their interactions. Continuous predictors were standardized (centered and scaled), and categorical variables were coded using sum contrasts. Statistical significance of predictors was assessed using Type III likelihood-ratio χ² tests implemented in the *car::Anova* function in R. A set of 37 candidate models representing different combinations of microsite attributes and interaction terms was evaluated (Table S1). These models allowed testing different hypotheses regarding the relative importance of soil properties and microsite type in determining seedling survival. The data shows that soil moisture is a critical driver of seedling survival. To further visualize seedling survival dynamics along the soil moisture gradient, soil moisture values were divided into quartiles, and survival proportions were calculated for each quartile by species and microsite. All statistical analyses were conducted in R Studio.

## 3.0 Results

### 3.1 Differences in soil attributes by microsite type

Soil moisture differed significantly between microsite types (Welch’s t-test: *t* = 8.68, *df* = 124.21, *p* < 0.001). Mean soil moisture was twice as high on intact ground microsites (0.270 ± 0.087 SD, *n* = 59) compared to mound tops (0.137 ± 0.088 SD, *n* = 72). Median moisture values were 0.265 for ground microsites and 0.118 for mound tops. Interquartile ranges did not overlap (ground: 0.210-0.303; top: 0.095-0.145), indicating consistently wetter ground soils and consistently drier mound-top soils (Fig. 2a). Soil organic matter (SOM) was also significantly higher with ground microsites (median = 0.066, IQR = 0.050) compared to mound tops (median = 0.033, IQR = 0.023; Wilcoxon rank-sum test: *W* = 3452, *p* < 0.001; Fig. 2b). In contrast, soil pH did not differ significantly between microsite types (Welch’s t-test: *t* = 0.968, *df* = 128.97, *p* = 0.335). Median pH values were 6.35 for ground microsites and 6.22 for mound tops (Fig. 2c). Soil texture differed between microsite types, with finer-textured soils more common on mound tops and coarser-textured soils more frequent with intact ground microsites (Fig. 2d).

**Fig. 2a.**
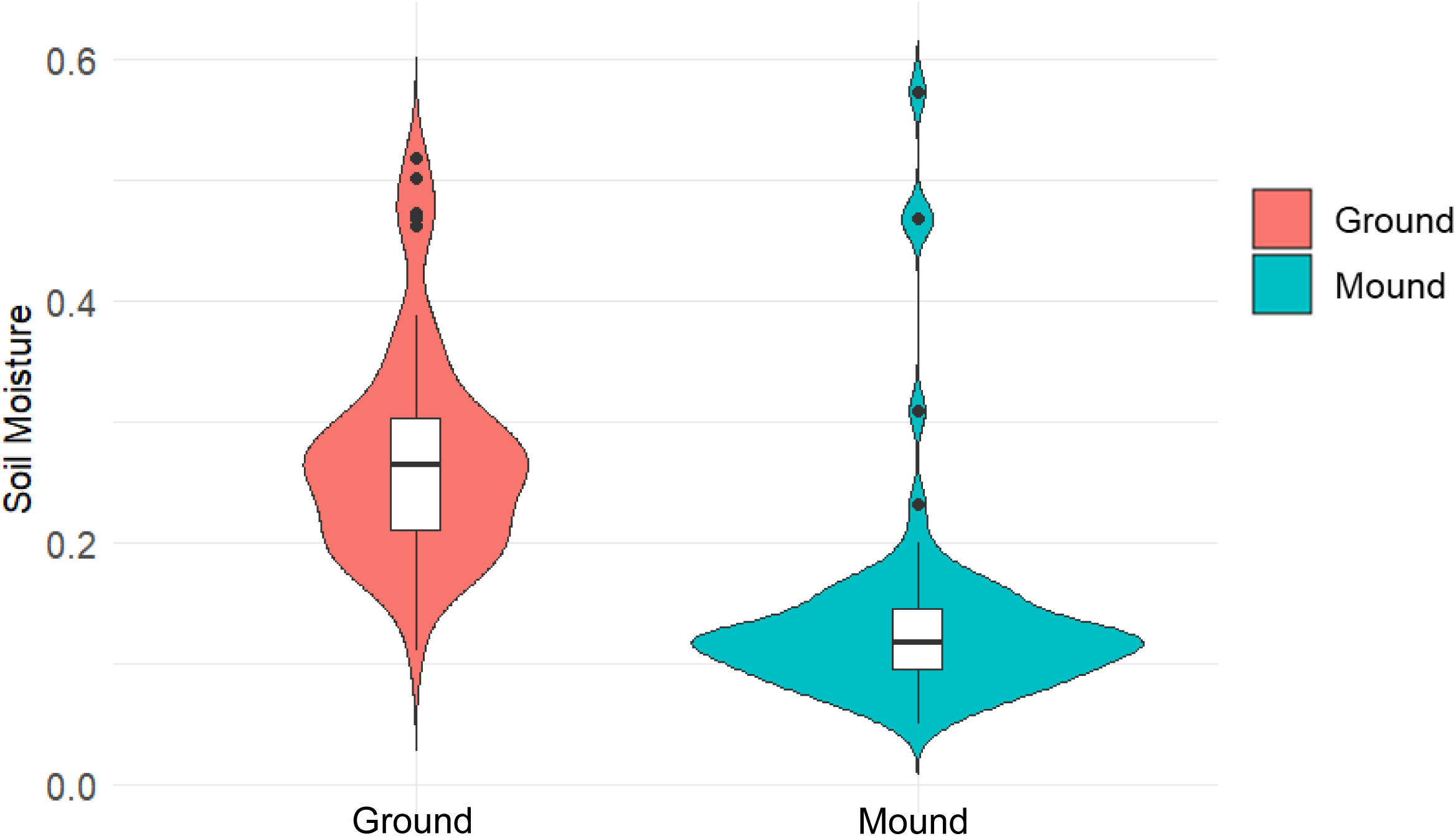
Soil moisture by microsite type

**Fig. 2b.**
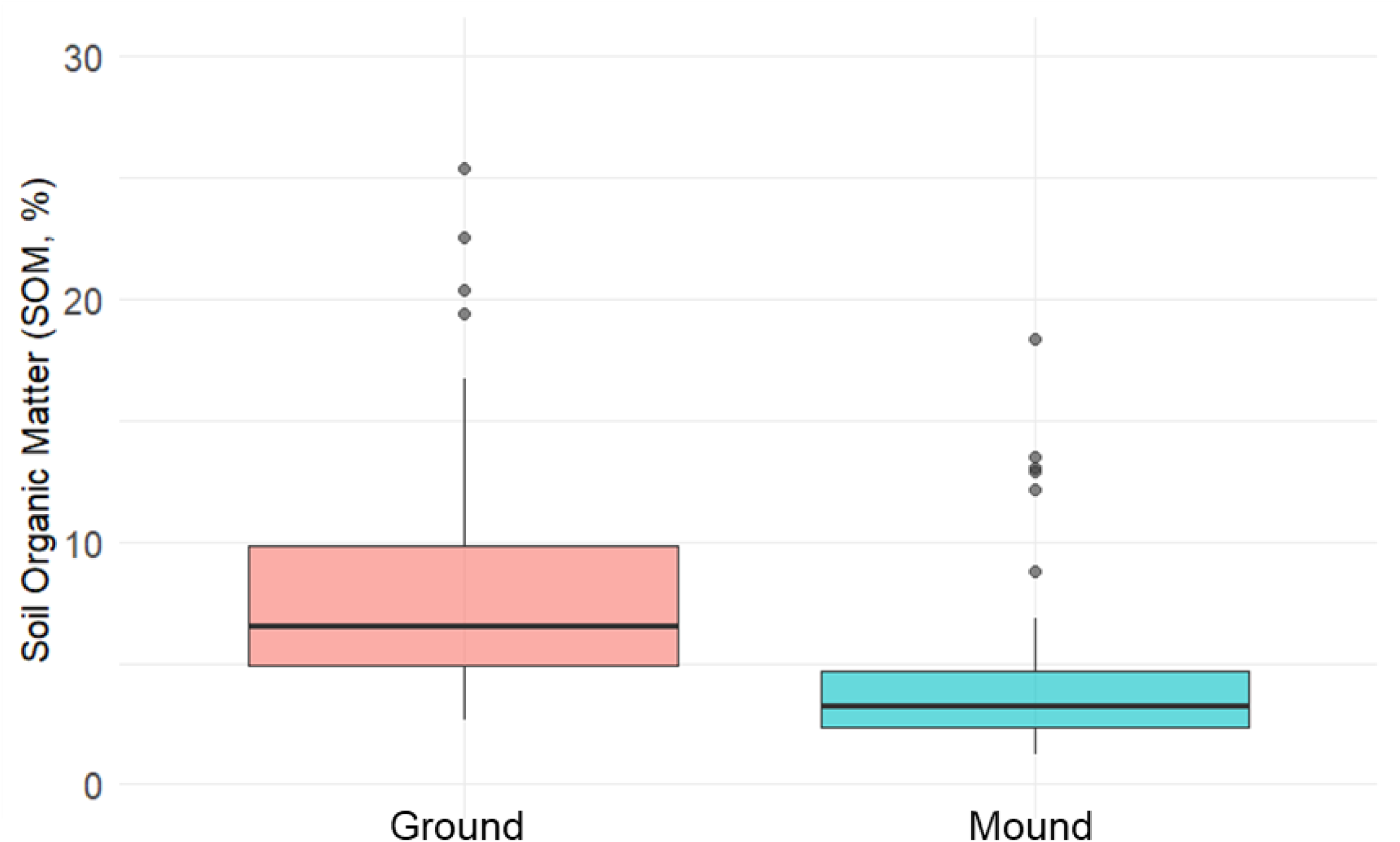
Soil organic matter (SOM) by microsite type

**Fig. 2c.**
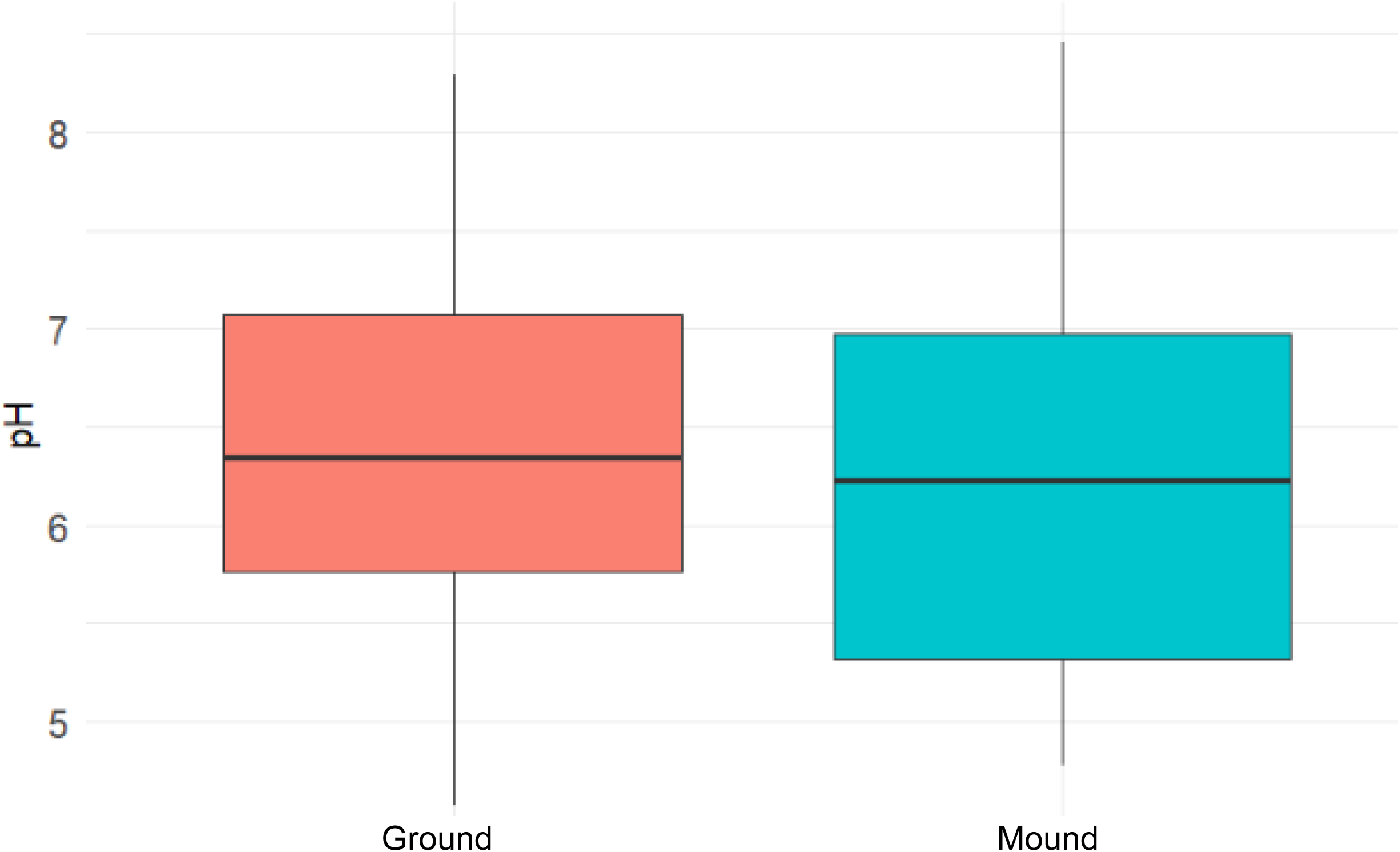
Soil pH by microsite type

**Fig. 2d.**
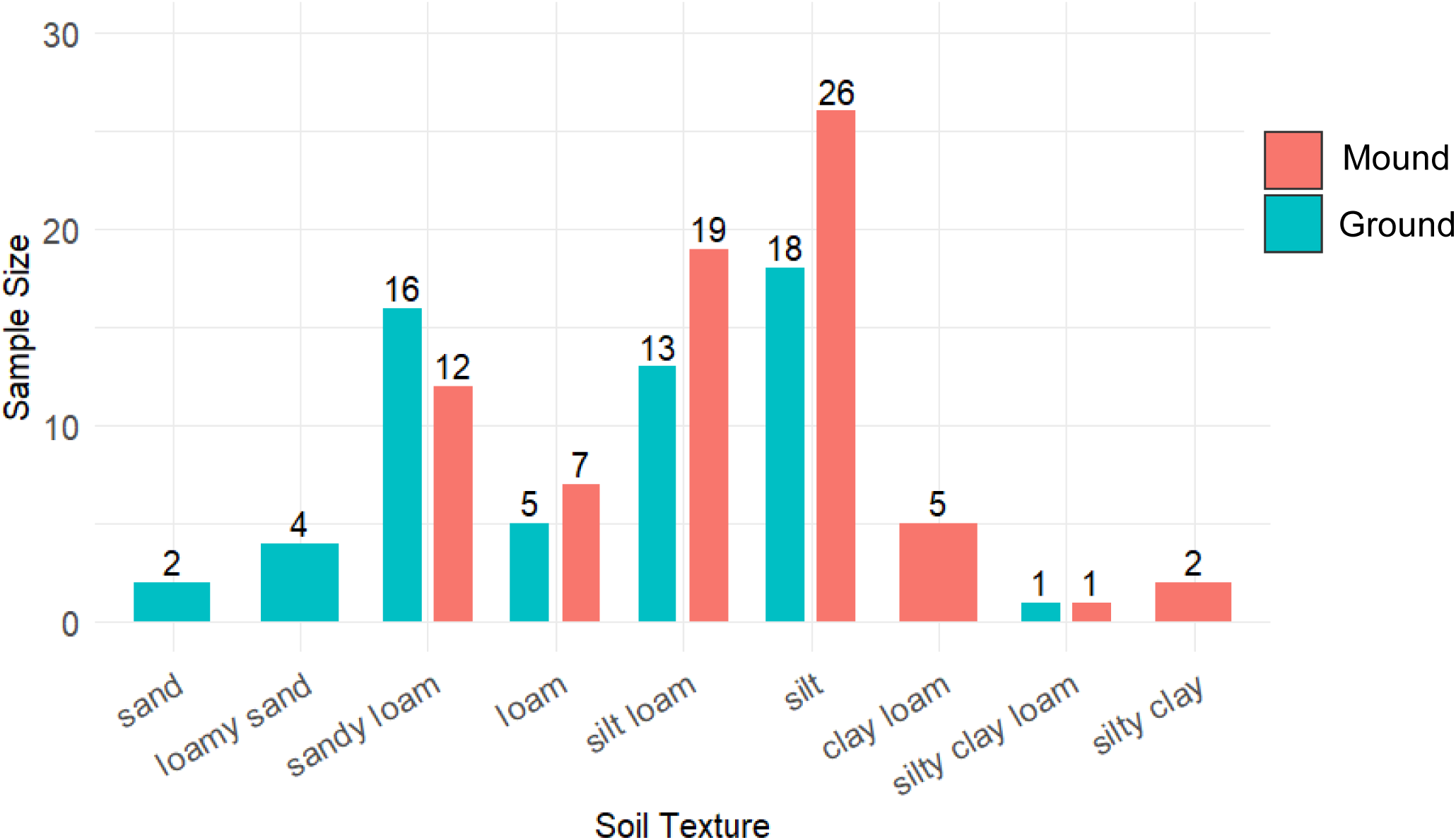
Distribution of soil texture by microsite

### 3.2 Seedling survival dynamics

Across both species combined, seedling survival was 39% on mound tops (*n=72*) and 41% on ground microsites (*n=59*) (Fig. 3a). Black cherry exhibited higher survival on mound tops (44%, *n* = 43) compared with intact ground microsites (28%, *n* = 29). In contrast, red oak survival was higher on intact ground microsites (53%, *n* = 30) compared to mound tops (31%, *n* = 29) (Fig. 3a). Seedling survival also varied slightly with the number of seedlings planted per mound. Mounds with one seedling had a mean survival rate of 37%, whereas mounds with two seedlings had a higher survival rate (50%) and mounds containing three or four seedlings had lowest survival rates of 33% and 25%, respectively.

**Fig. 3.**
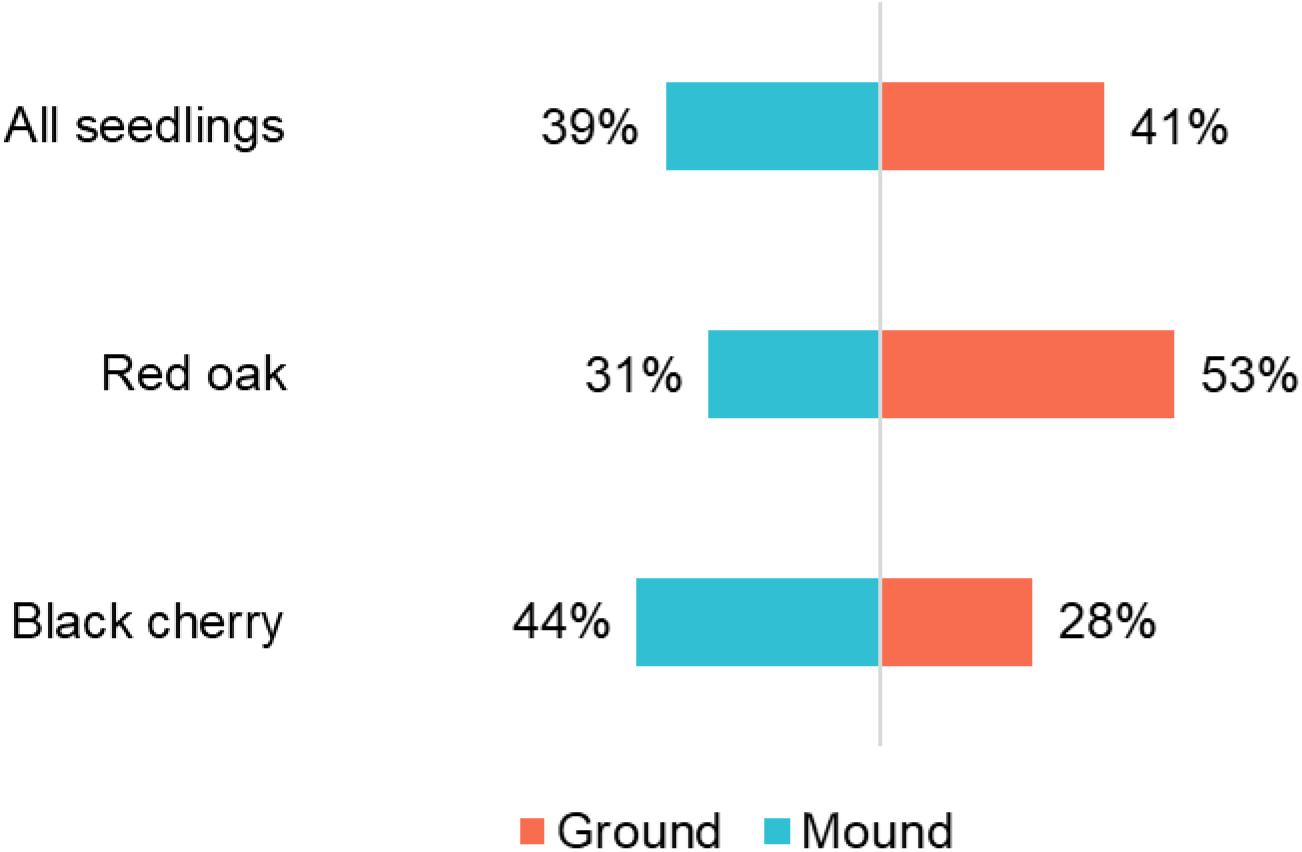
Seedling survival by species and microsite type (mound vs. ground).

### 3.3 Effects of microsite attributes on seedling survival

Generalized linear modeling indicated that soil moisture is a significant predictor of seedling survival (Species × Moisture, *p* = 0.019; Table S1). Red oak survival probability increased with increasing soil moisture, whereas black cherry survival probability decreased as soil moisture increased (Fig. 4a). These contrasting responses were evident across microsite types. Because windthrow mounds were consistently drier than intact ground microsites (Fig. 2a), predicted survival probabilities differed by microsites (Fig. 4b). Quartile analyses further illustrate these patterns. With intact ground microsites (Fig. 4c), black cherry survival was highest in the driest quartile (Q1: 56%) and declined in wetter quartiles (Q2: 0%, Q3: 29%, Q4: 17%). In contrast, red oak survival was lowest in the driest quartile (Q1: 17%) and increased in wetter quartiles (Q2: 50%, Q3: 75%, Q4: 62%). On windthrow mound microsites (Fig. 4d), black cherry survival increased with increasing moisture, rising from 27% in the driest quartile (Q1) to 60% in the wettest quartile (Q4). Red oak survival was lowest in the drier quartiles (Q1: 29%, Q2: 12%) and higher in wetter quartiles (Q3: 50%, Q4: 38%).

**Fig. 4a.**
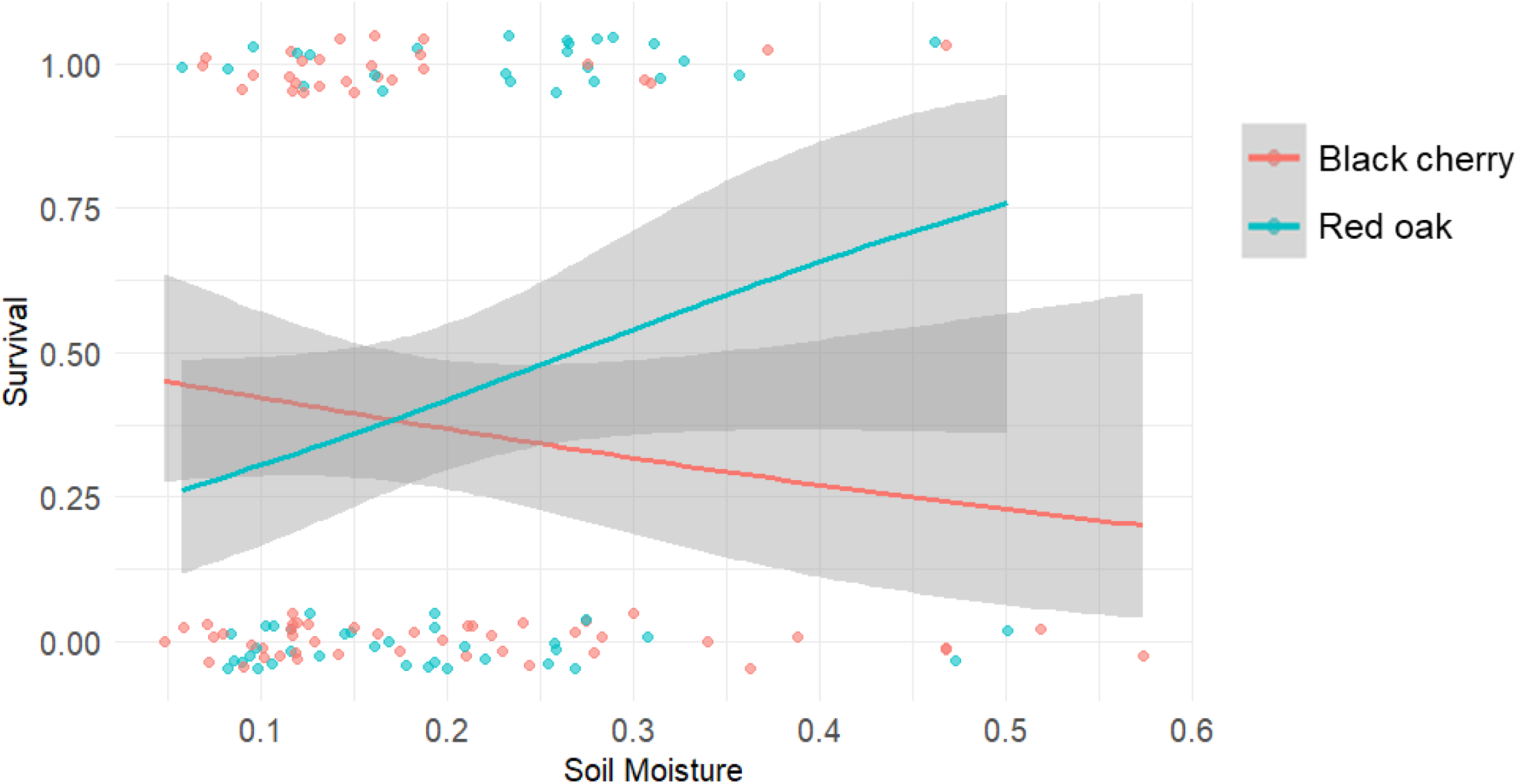
Seedling survival by species and soil moisture

**Fig. 4b.**
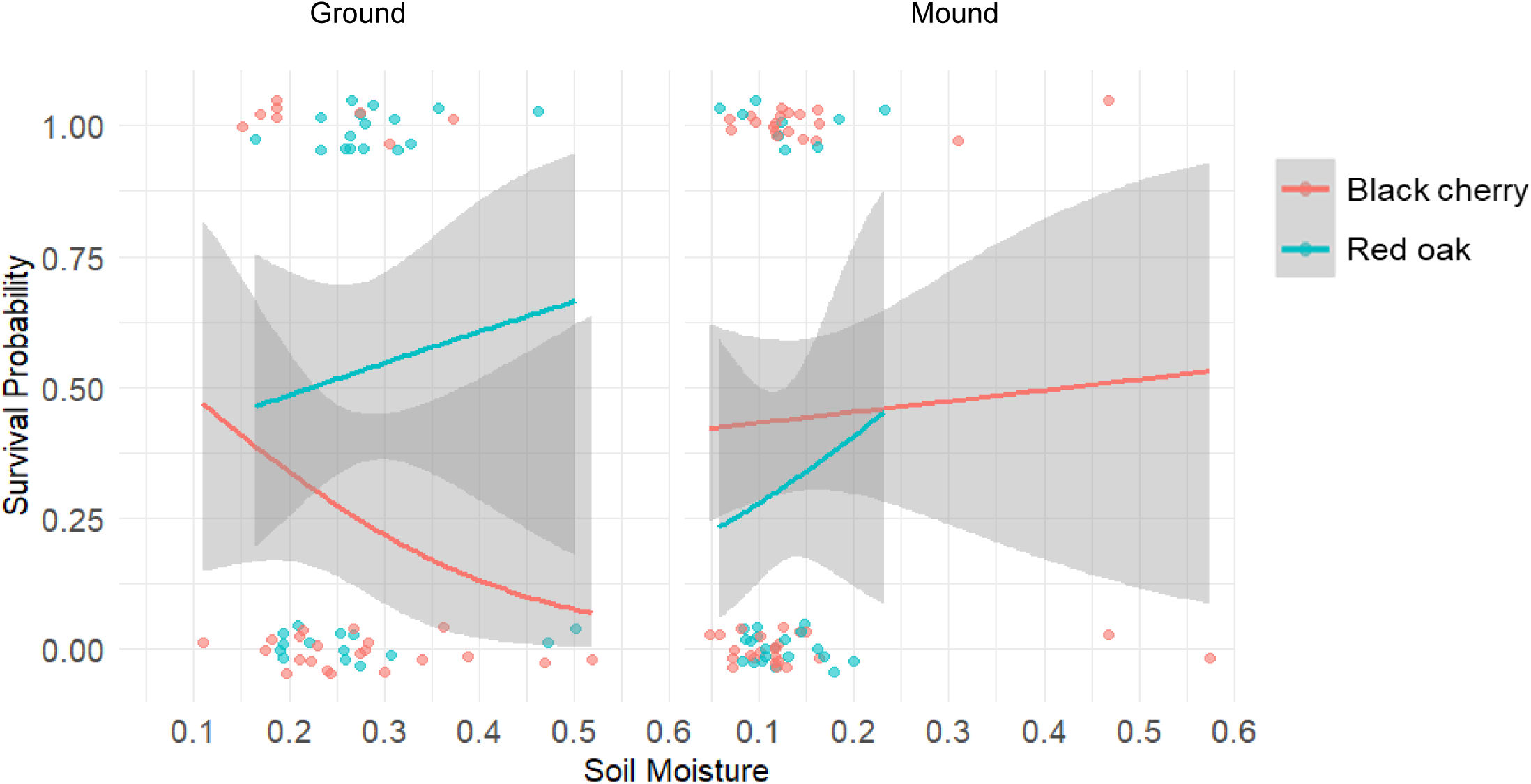
Seedling survival by species, soil moisture and microsite type.

**Fig. 4c.**
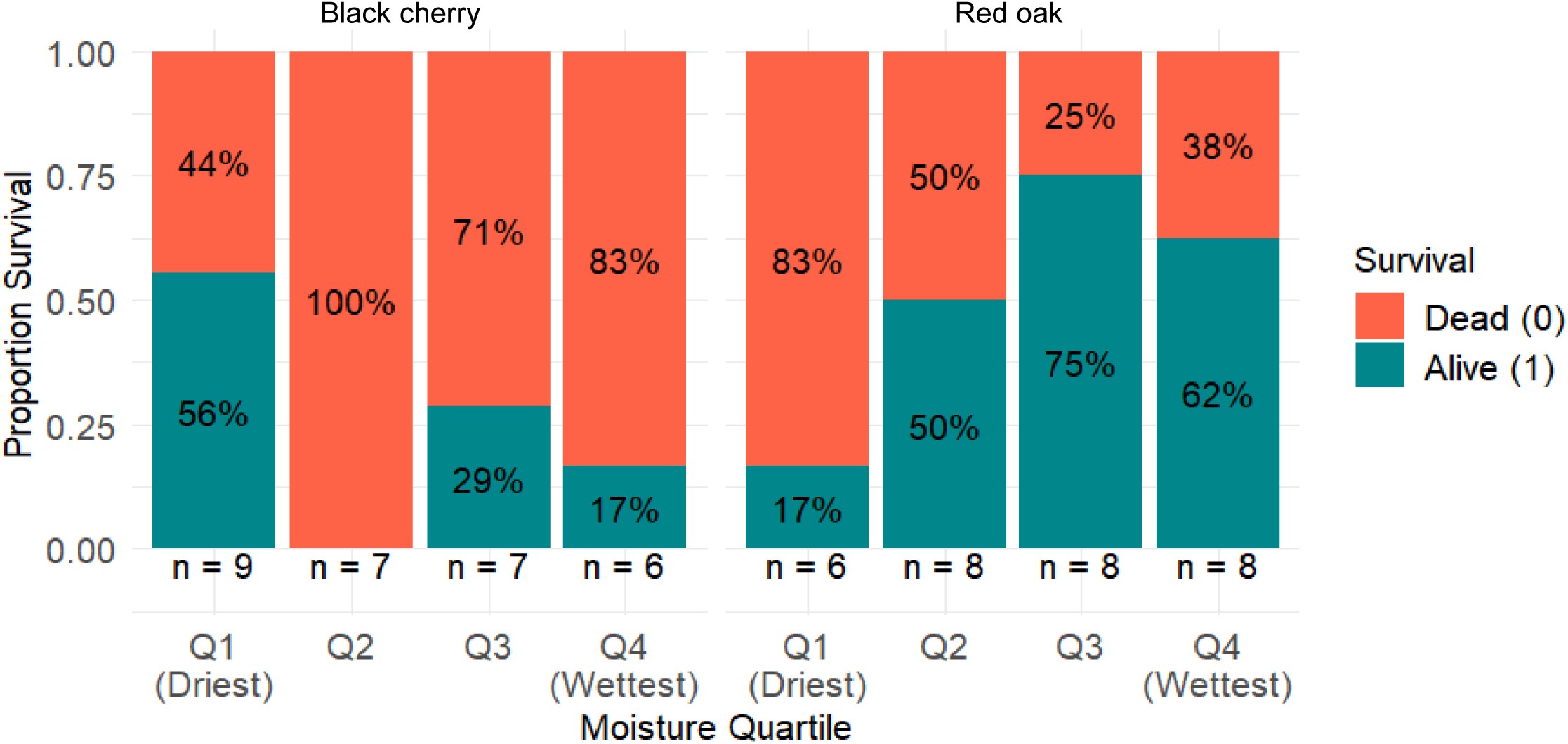
Seedling survival by species and moisture quartile for ground microsite type

**Fig. 4d.**
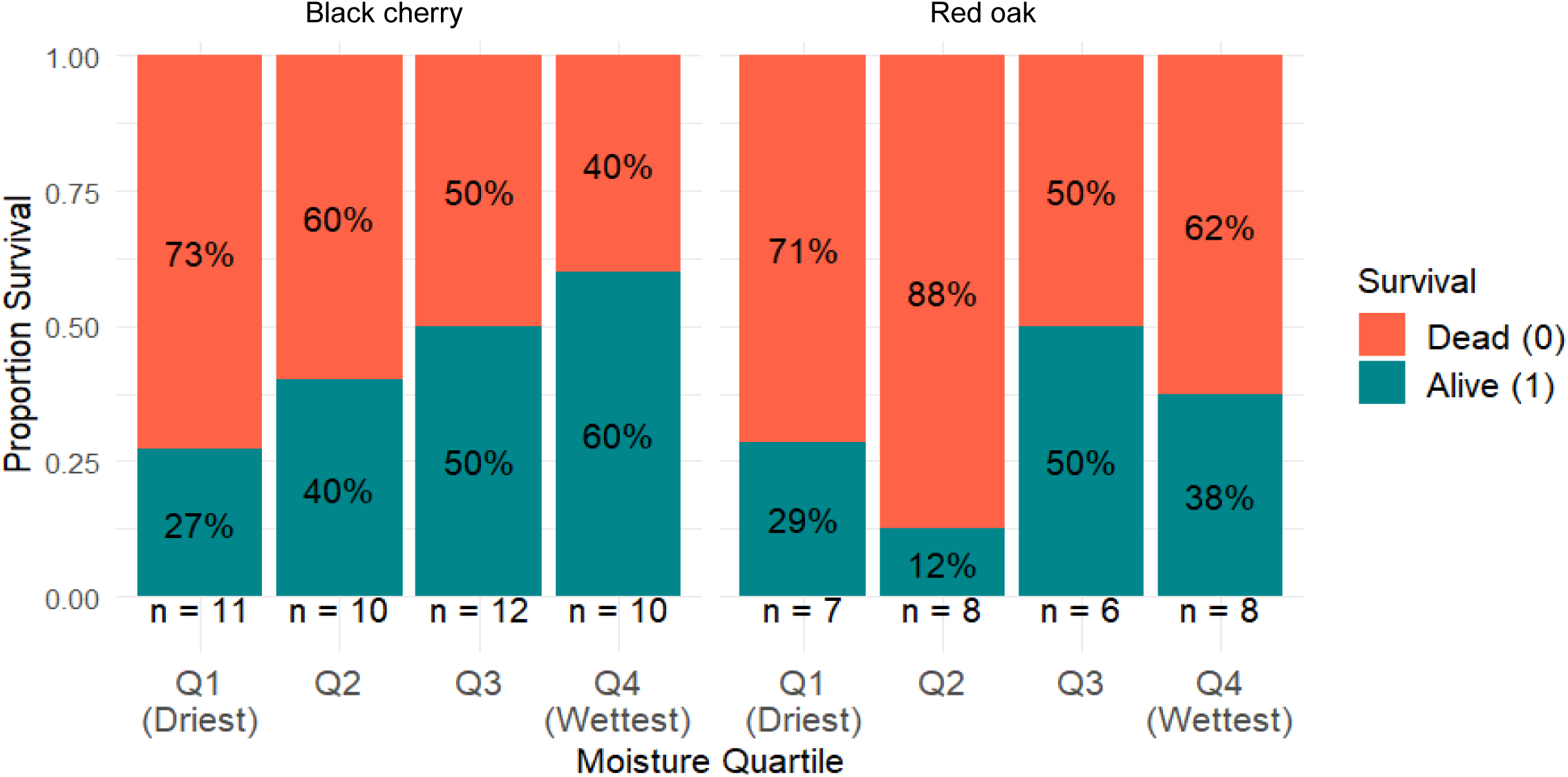
Seedling survival by species and moisture quartile for mound microsite type.

## 4.0 Discussion

The region of southern Ontario (Niagara escarpment) is designated as a UNESCO Biosphere reserve due to the rarity and richness of biodiversity. The region also has high ungulate densities (See sample photos and analysis of deer abundance in the supplementary section Fig. S2 - S7). This region also experiences high windstorm events (e.g. Munoz and Anyomi 2026), consequently, windthrow mounds are common within the conservation areas (Fig. 1). Windthrow generated mounds can serve as effective avenues for restoring deer over browsed species but a thorough understanding of the attributes of these microsites is necessary for successful integration into forest management planning and restoration activities. This study compared soil attributes of 84 windthrow generated microsites to those of adjacent intact ground microsites and evaluated their influence on the survival of red oak and black cherry seedlings in the old-growth Carolinian forest in Hamilton.

### 4.1 Moisture as the primary driver of microsite effects

Ground microsites consistently had higher soil organic matter (Fig. S1) than mound tops, exhibiting nearly twice the mean moisture content and a broader moisture range (Fig. 2a), than disturbed mound top microsite type. Red oak survival increased with increasing moisture, consistent with its known tolerance of moist soils (e.g. Sander, 1990; Kuehne et al., 2014; Nosko et al., 2022). In contrast, black cherry survival declined as moisture increased on ground microsites, suggesting potential sensitivity to oxygen limitation or waterlogging in compacted soils. Quartile analyses further illustrate these dynamics. On ground microsites (Fig. 4c), black cherry-often described as drought-tolerant (Barkley, 2007) achieved its highest survival in the driest soils (Q1, 56%), whereas red oak survival in these soils was only 17%, indicating vulnerability to desiccation stress. Oak survival increased substantially in wetter ground soils, reaching up to 71% in Q3, while cherry survival declined under these wetter conditions. On disturbed sites (mound tops), however, black cherry survival increased with moisture, reaching 60% in the wettest quartile (Q4). This pattern suggests that moisture itself is not inherently detrimental to cherry seedlings. Rather, cherry appears sensitive to oxygen limitation associated with poorly aerated soils, a condition common on the compacted forest floor. Physiological studies indicate that oxygen limitation under waterlogged conditions can impair black cherry performance (Wang et al., 2023). Mounds provide well-drained and aerated soils, allowing oxygen to remain available in the rooting zone even when moisture increases. Under these conditions, cherry seedlings can benefit from both water availability and adequate aeration. Red oak, meanwhile, maintained relatively low survival (≤50%) across all moisture levels on disturbed sites (mound tops). Because mound tops are generally drier than ground microsites, they appear to not have the optimum moisture conditions that favor oak survival. The summer of 2025 was typically warm and may have contributed to the contrast between red oak and black cherry regarding the sensitivity to moisture, highlighting the mediating role of recently loosened and less compacted soil in successful seedling establishment.

### 4.2 Life-history strategies and species responses to tip-up mounds

Red oak and black cherry represent contrasting strategies that may have influenced their responses to microsite conditions. Oak seedlings grow slowly and allocate early resources toward deep and robust root systems rather than rapid above-ground growth. This strategy allows oak to persist in shaded understories as part of advanced regeneration (Dillaway and Stringer 2006, Nosko et al., 2022), where seedlings remain suppressed until canopy openings permit accelerated growth. Black cherry on the other hand is a fast-growing, shade-intolerant species that relies on rapid establishment following disturbance (Marquis, 1990). Its fine root system requires well-aerated soils, and its growth is closely tied to light availability. Consequently, cherry performs poorly on compacted and shaded intact ground soils but had higher survival in disturbed microsites that provide increased light and along an increasing gradient of soil moisture. Although black cherry is sometimes described as drought-tolerant, its apparent advantage in drier soils may partly reflect avoidance of oxygen limitation in compacted ground soils rather than a direct preference for drought conditions. Tip-up mounds expose loose mineral soil, reduce litter layers, and open canopy gaps, creating conditions that closely match the ecological requirements of this disturbance-adapted species.

### 4.3 Windthrow generated mounds as ecological filters

Stand development and ecological succession lead to canopy closure, and earthworm invasion in many north American soils have been associated with increased soil compaction, high moisture retention, and reduced aeration in many northern forests (Frelich et al., 2006; Tsogbadrakh et al., 2024). These changes may lead to chronically low oxygen availability in the rooting zone, particularly in undisturbed ground microsites.

For species such as black cherry that require well-drained and aerated soils, these conditions may limit successful establishment. Similar patterns have been documented across eastern North America, where mesic species have increased while disturbance-adapted or drought-tolerant species have declined (Woodbridge et al., 2022). In this context, forest floors function as long-term environmental filters that shape regeneration patterns and influence future forest composition in the absence of disturbances.

Our findings suggest that windthrow generated mounds function as ecological filters (Keddy, 1992) that selectively favor species with certain functional traits. Mounds modify several environmental factors simultaneously: they increase light availability, improve soil aeration through loosened soil structure, and enhance drainage due to their elevated position (Lutz 1940; Yoshida, 2021). These changes alleviate many of the constraints faced by fast growing shade intolerant species such as black cherry while simultaneously removing the stable moisture conditions that shade tolerant species such as oak are adapted to.

## 5.0 Conclusion

The results indicate that within the Carolinian forest of southern Ontario, windthrow generated mounds favor species that require well-drained and aerated soils, such as black cherry, whereas the surrounding intact forest floor characterized by higher soil organic matter, higher soil moisture, greater soil compaction, and persistent shading favors species tolerant of these conditions, such as red oak. These contrasting niches illustrate small-scale disturbances such as tree uprooting impact local forest microsite conditions and seedling establishment. From ecological restoration perspective, these findings challenge the common assumption that planting directly into undisturbed forest floor soils is universally suitable for seedling establishment.

## Supporting information

Duan and Anyomi_2026_Supple Documents

## Acknowledgements

The project was funded by TD Friends of the Environment grant (ID: 94206595), Redeemer University Internal Research grant (ID: 2025-404-IRG-08) and MITACS GRI award (ID: 46064). We are grateful to Jessica Hart for helping to find the mounds used in this study. We are also grateful to the Hamilton Conservation Authority for allowing this work to be conducted in the conservation areas in Hamilton. We are also thankful to McMaster University for allowing us to use mounds in the McMaster Forest Preserve for the study. We are also grateful to the Bruce Trail Conservancy for allowing us to use mounds along the Bruce trail by Tiffany Falls for this study.

