## Supplementary material for "Windthrow-generated mounds create contrasting regeneration niches for red oak and black cherry in a deer-browsed Carolinian forest": Duan and Anyomi_2026_Supple Documents

SUPPLEMENTARY TABLE

Table S1. Seedling survival candidate models

| **Model** | **k** | **AIC** | **DeltaAIC** | **BIC** |
| --- | --- | --- | --- | --- |
| Survival ~ Moisture_c:Species | 3 | 177.6133 | 0 | 186.2389 |
| Survival ~ Species:Position | 5 | 178.5693 | 0.956021 | 190.0701 |
| Survival ~ Species * Position | 4 | 178.5693 | 0.956021 | 190.0701 |
| Survival ~ pH_c | 2 | 178.9725 | 1.359179 | 184.7229 |
| Survival ~ Moisture_c:Species:Position | 5 | 179.3041 | 1.690823 | 193.6801 |
| Survival ~ Moisture_c * Species | 4 | 179.3651 | 1.751825 | 190.8659 |
| Survival ~ SOM_c | 2 | 179.6418 | 2.0285 | 185.3922 |
| Survival ~ Species | 2 | 179.6782 | 2.064919 | 185.4286 |
| Survival ~ Moisture_c + Species + Position + pH_c + SOM_c + Moisture_c:Species + Moisture_c:Position | 8 | 179.7747 | 2.161396 | 202.7763 |
| Survival ~ Moisture_c | 2 | 179.8745 | 2.261192 | 185.6249 |
| Survival ~ Position | 2 | 179.9563 | 2.342952 | 185.7066 |
| Survival ~ pH_c + Species | 3 | 180.755 | 3.141748 | 189.3806 |
| Survival ~ Moisture_c + Species + Position + pH_c + SOM_c + Moisture_c:Position + Species:Position | 8 | 180.8008 | 3.187541 | 203.8024 |
| Survival ~ pH_c:Species | 3 | 180.8339 | 3.220649 | 189.4595 |
| Survival ~ SOM_c + Species | 3 | 181.3647 | 3.75137 | 189.9903 |
| Survival ~ Moisture_c + Species | 3 | 181.5731 | 3.959837 | 190.1987 |
| Survival ~ Moisture_c:Position | 3 | 181.6032 | 3.98986 | 190.2288 |
| Survival ~ SOM_c:Species | 3 | 181.6172 | 4.003898 | 190.2428 |
| Survival ~ Species + Position | 3 | 181.6559 | 4.042646 | 190.2815 |
| Survival ~ Moisture_c + Position | 3 | 181.8744 | 4.261143 | 190.5 |
| Survival ~ Moisture_c + Species + Position + pH_c + SOM_c + Species:Position | 7 | 182.284 | 4.670703 | 202.4104 |
| Survival ~ pH_c * Species | 4 | 182.5885 | 4.975212 | 194.0893 |
| Survival ~ Moisture_c + Species + Position + pH_c + SOM_c + Moisture_c:Species | 7 | 183.2554 | 5.642088 | 203.3818 |
| Survival ~ Moisture_c * Species + Position + pH_c + SOM_c | 7 | 183.2554 | 5.642088 | 203.3818 |
| Survival ~ Moisture_c + Species + Position + pH_c + SOM_c + Moisture_c:Position | 7 | 183.2682 | 5.654891 | 203.3946 |
| Survival ~ Moisture_c * Position + Species + pH_c + SOM_c | 7 | 183.2682 | 5.654891 | 203.3946 |
| Survival ~ SOM_c * Species | 4 | 183.3158 | 5.702494 | 194.8166 |
| Survival ~ Moisture_c + Species + Position | 4 | 183.5697 | 5.956352 | 195.0704 |
| Survival ~ Moisture_c * Position | 4 | 183.5991 | 5.985809 | 195.0999 |
| Survival ~ Moisture_c + Species + Position + pH_c + SOM_c + Moisture_c:Species + Species:Position | 8 | 184.0397 | 6.426412 | 207.0413 |
| Survival ~ Moisture_c * Species * Position | 8 | 184.4637 | 6.850412 | 207.4653 |
| Survival ~ Moisture_c + Species + Position + pH_c + SOM_c | 6 | 184.8499 | 7.236574 | 202.1011 |
| Survival ~ Moisture_c + Species + Position + pH_c + SOM_c + Moisture_c:Species + Moisture_c:Position + Soil.Texture | 16 | 191.0244 | 13.41114 | 237.0276 |
| Survival ~ Moisture_c + Species + Position + pH_c + SOM_c + Species:Position + Soil.Texture | 15 | 191.9977 | 14.38441 | 235.1257 |
| Survival ~ Moisture_c + Species + Position + pH_c + SOM_c + Moisture_c:Species + Soil.Texture | 15 | 194.0934 | 16.48015 | 237.2214 |
| Survival ~ Moisture_c + Species + Position + pH_c + SOM_c + Moisture_c:Position + Soil.Texture | 15 | 194.1215 | 16.50819 | 237.2494 |
| Survival ~ Moisture_c + Species + Position + pH_c + SOM_c + Soil.Texture | 14 | 195.3754 | 17.7621 | 235.6282 |

SUPPLEMENATRY FIGURES

***Inter-relationship between soil attributes***

Soil moisture increased nonlinearly with soil organic matter (SOM), explaining 53.5% of the variance in moisture (r=0.665, Figure S1a). This suggests that higher organic matter content contributes to increased water retention in ground microsites. Soil moisture showed a weak but significant negative correlation with soil texture fineness (Spearman’s ρ = −0.18, p = 0.040; Figure S1b). Coarser soils were associated with higher moisture values. This relationship likely reflects the co-variation between soil texture and SOM across microsites rather than a direct causal relationship between texture and moisture. Soil pH showed a weak but significant positive relationship with soil moisture (β = 0.021, R² = 0.032, p = 0.039; Figure S1c).

| 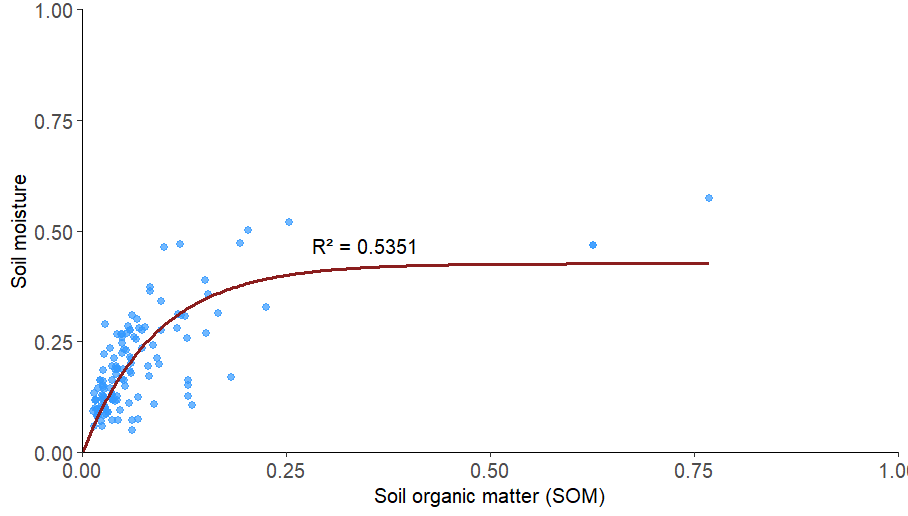  **a** | 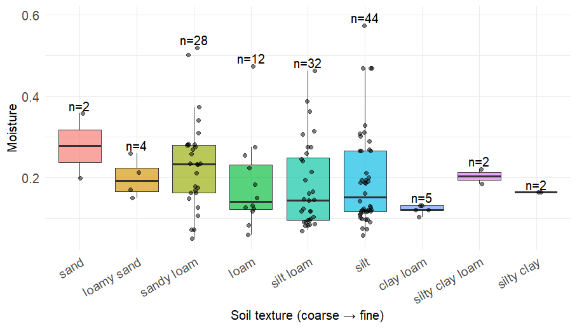  **b** |
| --- | --- |
| 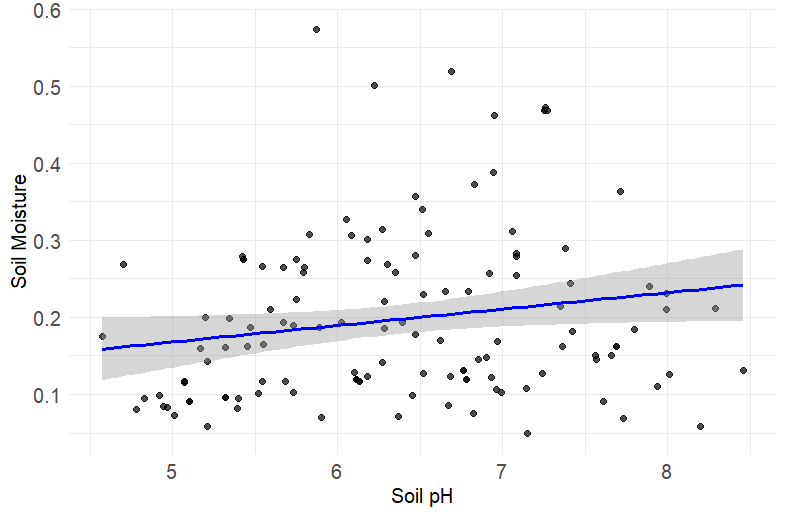  **c** |  |

Figure S1. Variation in soil moisture by a) soil organic matter (SOM), b) soil texture, and c) soil pH.

*Trail Camera Monitoring*
To document wildlife presence and white-tailed deer browsing activities, 10-Browning Strike Force FHDR motion-triggered trail cameras were installed next to the mounds. Cameras were positioned with the mound as the primary focal point, while also including as much of the adjacent ground area as possible to capture a broader range of wildlife activity. This setup maximized the likelihood of detecting potential herbivores, while also documenting other fauna present in the vicinity. Camera was monitored every 2-weeks. Trail camera data covered June 20^th^ to September 5^th^ 2025 and were reviewed manually and with assistance from Grok 4 series AI model (xAI, 2025). Species richness was calculated as the number of unique species in an area.

Wildlife species richness ranged from 2 to 7 across conservation area and the number of white-tailed deer detected ranged from 11 to 385. Conservation areas with relatively low wildlife species richness recorded higher white-tailed deer numbers (rho=-0.66, p>0.05). White-tailed deer observed has various fur coloration and pattern (Figure S3). A young white-tailed deer is often seen with an adult browsing typically during the day (Figure S4). A mature male deer (buck) is typically seen browsing at night alone (Figure S5). Coyotes and racoons were seen mostly at night across the conservation areas (Figure S6). Wild turkeys and some bird species were also observed by the mounds during the day (Figure S7).

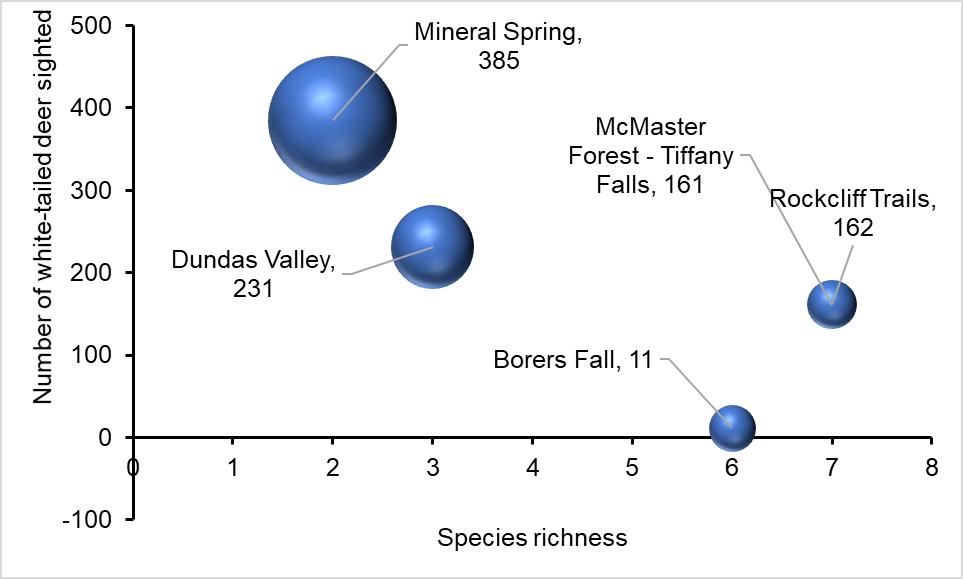

Figure S2. Number of white-tailed deer (detected by trail camera) by the total wildlife species richness. The size of the bubble refers to the duration of monitoring which ranged from 20 to 56 days (June - September 2025).

| 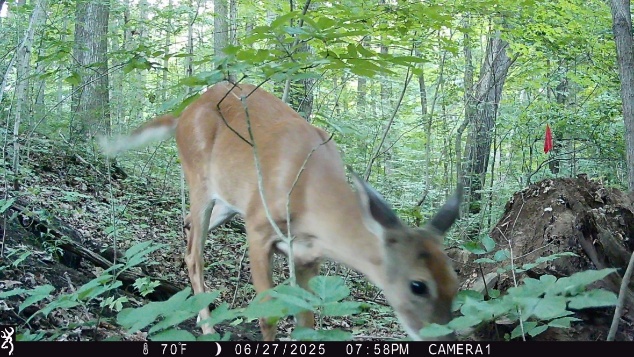  a | 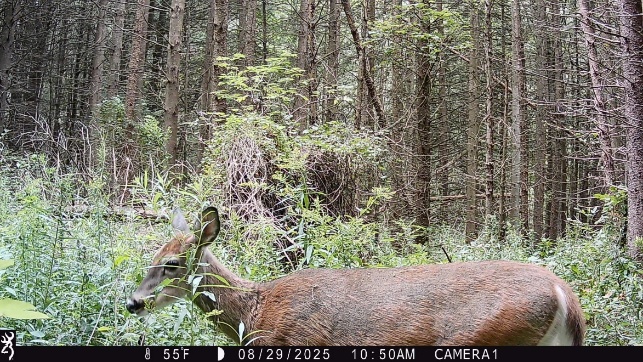  b |
| --- | --- |
| 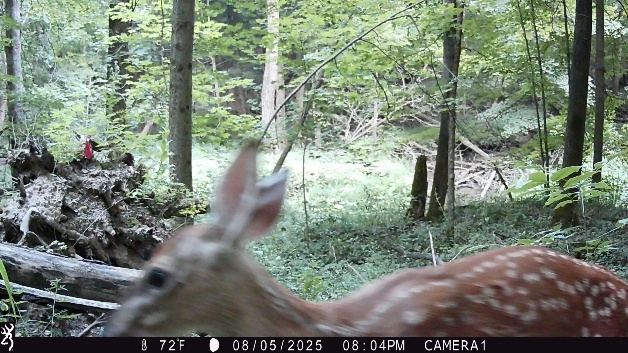  c | 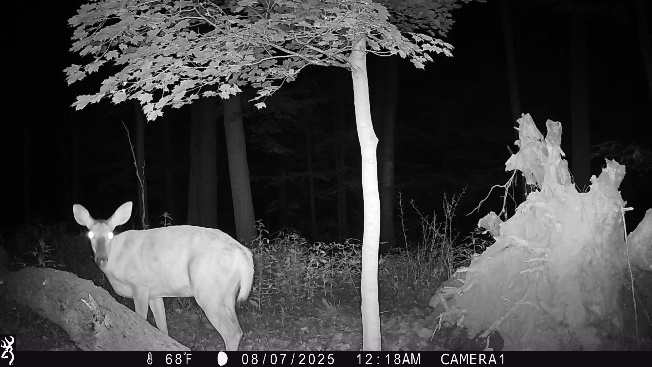  d |

Figure S3 a) smooth shiny brown fur of white-tailed deer browsing at Borer’s fall conservation area, Hamilton, Ontario. b) A reddish-brown fur of deer (*Odocoileus virginianus*) detected at the Mineral spring area, Hamilton, Ontario. c) White-tailed deer with smooth reddish-brown fur and white spots detected at the Dundas valley conservation area. d) White-tailed deer at McMaster Research forest-Tiffany falls conservation area suspected to be leucistic white-tailed deer (appearance may be due to night vision)

| 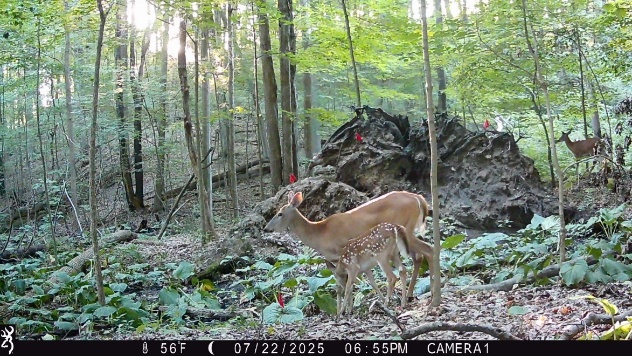  a | 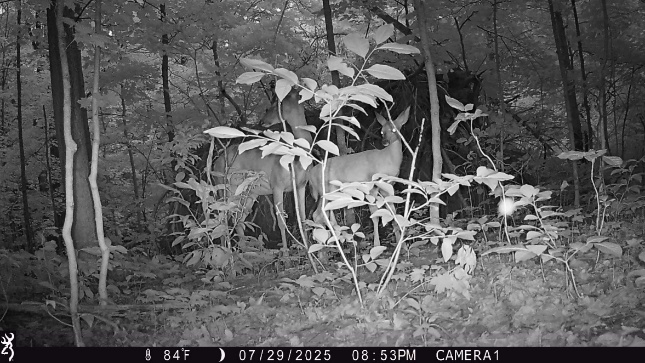  b |
| --- | --- |
| 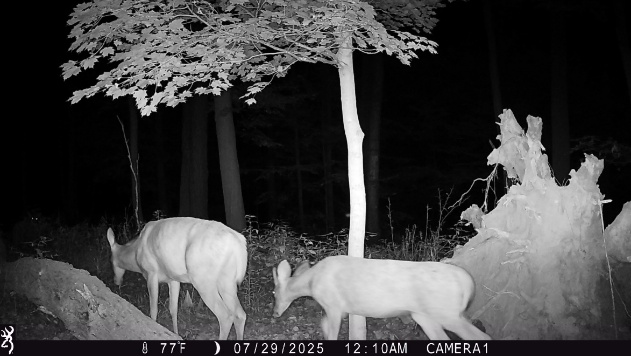  c | 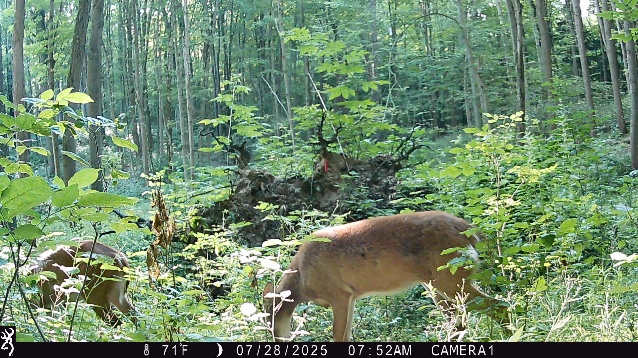  d |

Figure S4 a) A young deer and likely mother (*Odocoileus virginianus*) detected at the Mineral spring area, Hamilton, Ontario. b) White-tailed deer (*Odocoileus virginianus*), likely mother and young one detected at the Rockcliff trails area, Hamilton, Ontario. c) A young and adult white-tailed deer (*Odocoileus virginianus*) detected at the McMaster Research Forest (Tiffany Falls Conservation area), Hamilton, Ontario. D) A young and adult white-tailed deer (*Odocoileus virginianus*) detected at the Dundas valley conservation area, Hamilton, Ontario.

| 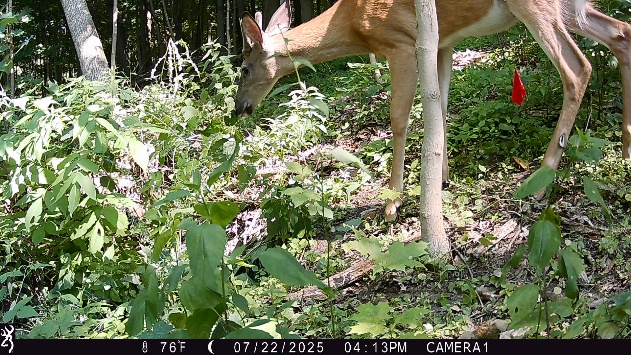  a | 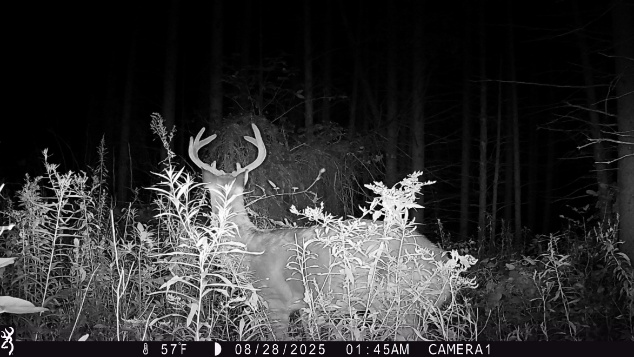  b |
| --- | --- |

Figure S5 a) A young buck browsing at the Tiffany Falls conservation area in Hamilton, Ontario. b) A mature buck (*Odocoileus virginianus*) detected at the Mineral spring area, Hamilton, Ontario.

| 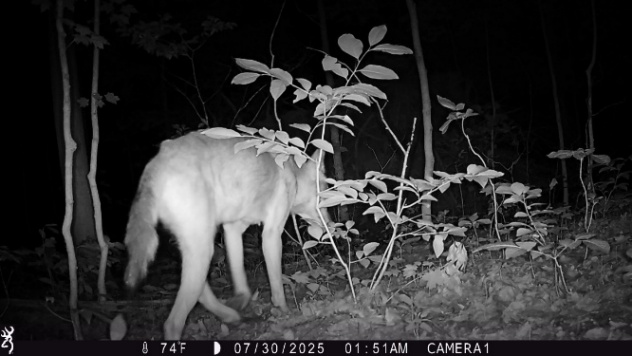  a | 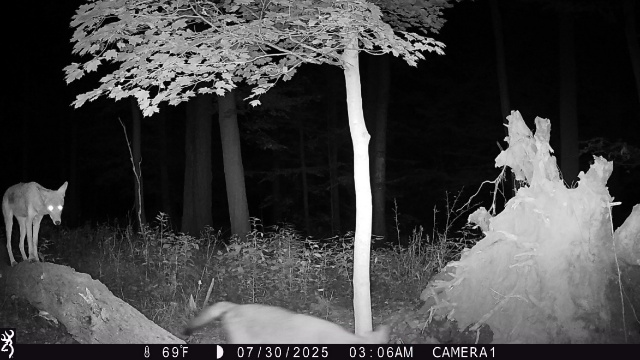  b |
| --- | --- |
| 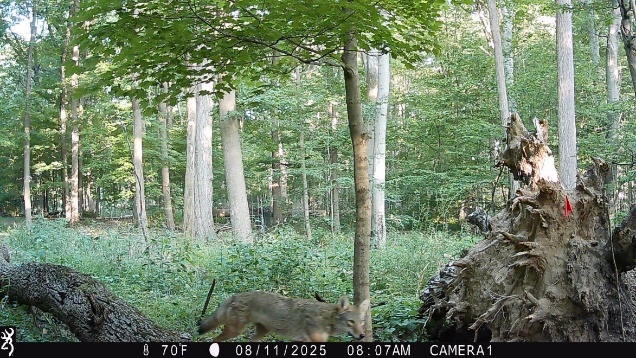  c | 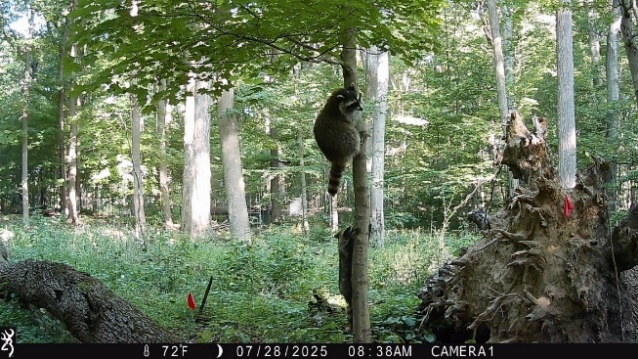  d |
| 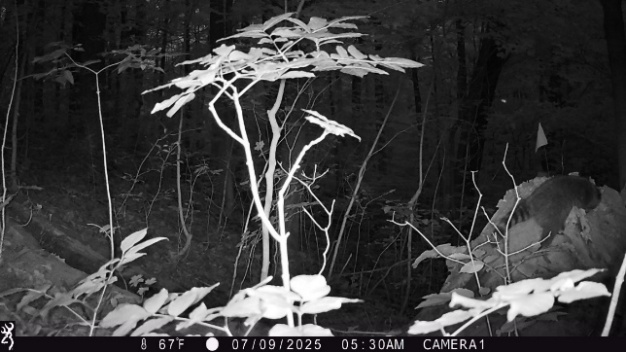  e | 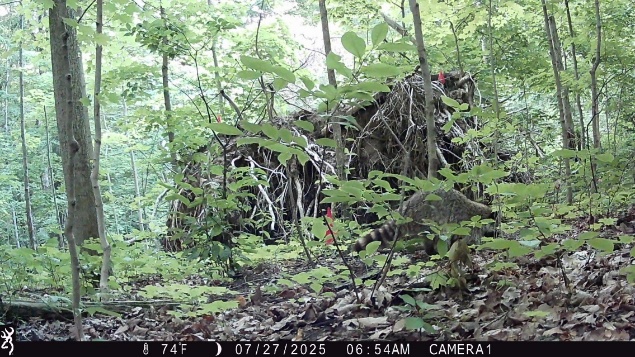  f |

Figure S6 a) Coyote (*Canis latrans*) detected at the Rockcliff trails area, Hamilton, Ontario, b) two coyotes (*Canis latrans*) detected at the McMaster Research Forest (Tiffany Falls Conservation area), Hamilton, Ontario, c) a coyote (*Canis latrans*) detected during the day at the McMaster Research Forest (Tiffany Falls Conservation area), Hamilton, Ontario. d) two racoons (*Procyon lotor*) detected at the McMaster Research Forest (Tiffany Falls Conservation area), Hamilton, Ontario, e) Raccoon (*Procyon lotor*) detected at the Borer’s fall conservation area, Hamilton, Ontario, f) Raccoon (*Procyon lotor*) detected at the Rockcliff trails area, Hamilton, Ontario

| 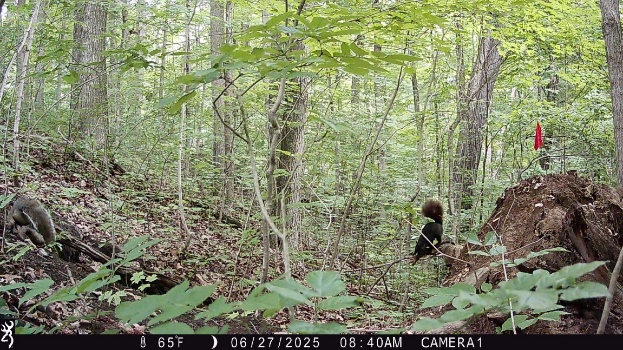  a | 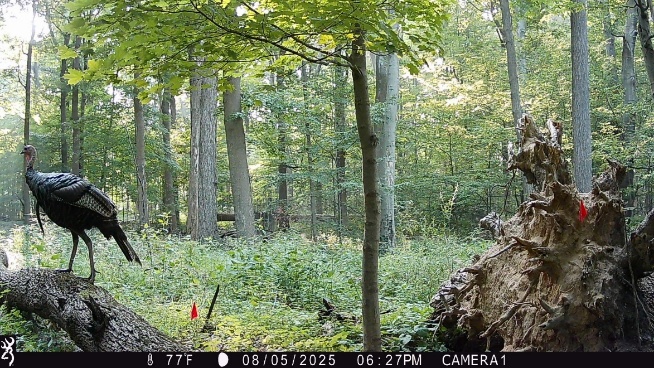  b |
| --- | --- |
| 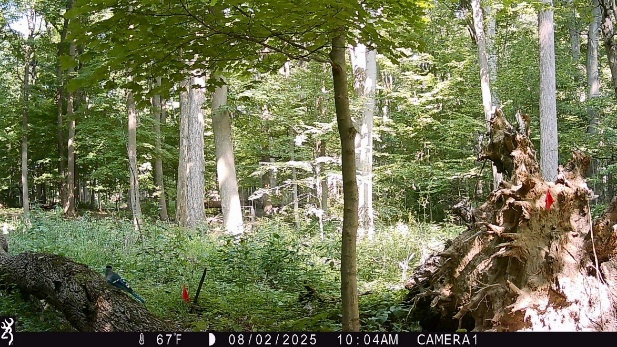  c | 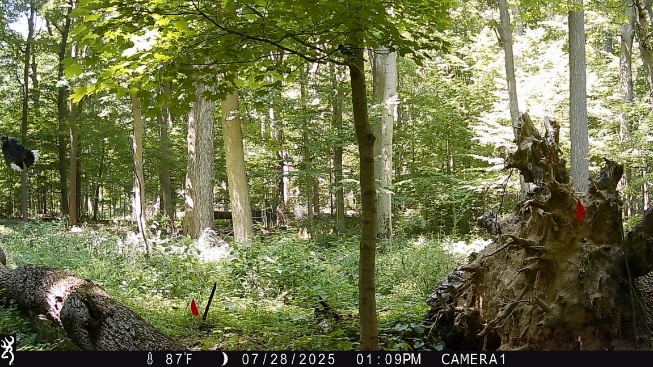  d |
| 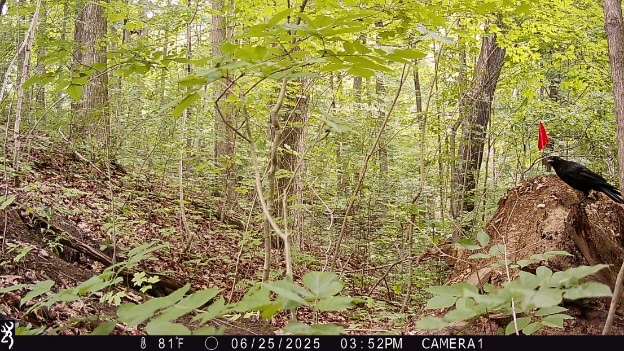  e | 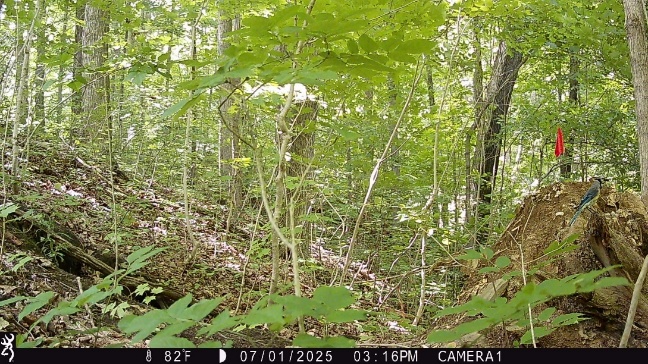  f |
| 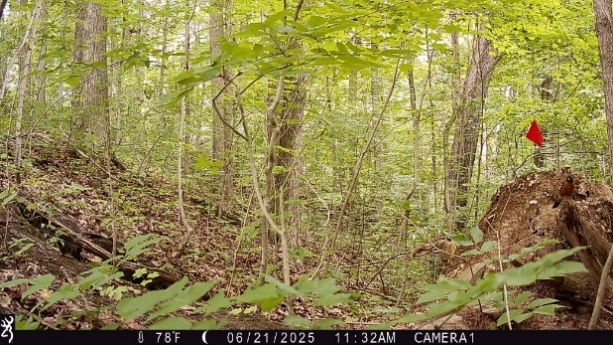  g | 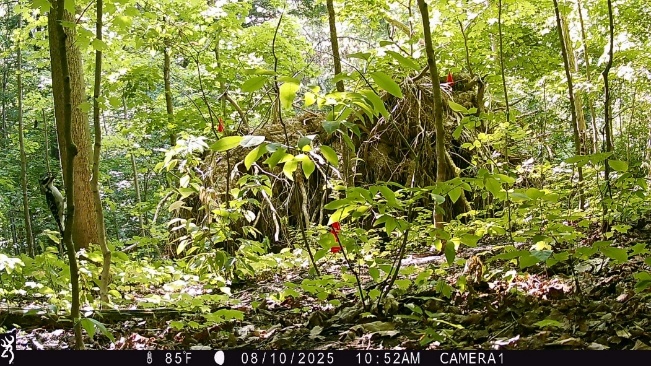  h |

Figure S7 a) eastern gray squirrel of both black and grey coloration (*Sciurus carolinensis*) were observed everywhere. b) Borer’s fall conservation area, Hamilton, Ontario. A wild turkey (*Meleagris gallopavo*) detected at the McMaster Research Forest (Tiffany Falls Conservation area), Hamilton, Ontario, c) a downy wood pecker *(Dryobates pubescens*) detected at the McMaster Research Forest (Tiffany Falls Conservation area), Hamilton, Ontario, d) a bird likely eastern king bird (*Tyrannus tyrannus*) detected at the McMaster Research Forest (Tiffany Falls Conservation area), Hamilton, Ontario, e) American crow (*Corvus brachyrhynchos*) detected at the Borer’s fall conservation area, Hamilton, Ontario. f) Blue jay (*Cyanocitta cristata*) detected at the Borer’s fall conservation area, Hamilton, Ontario, g) American robin (*Turdus migratorius*) detected at the Borer’s fall conservation area, Hamilton, Ontario, h) a downy wood pecker *(Dryobates pubescens*) detected at the Rockcliff trails area, Hamilton, Ontario. Please note that the quality of photos is not the best for accurate identification of bird species.
